# Effects of hCG or GnRH Treatment on Embryonic Mortality and Reproductive Performance in Buchi sheep of Cholistan Desert, Pakistan

**DOI:** 10.1101/2025.05.29.656925

**Authors:** Mubarra Anam, Mushtaq Hussain Lashari, Muhammad Saleem Akhtar, Mazhar Abbas

## Abstract

The present study was conducted on Buchi sheep at Government livestock farm, Jugaitpeer, district Bahawalpur. The objectives of this study were to investigates the genetic potential, birth and adult body weights, growth and turnover rates, and seasonal reproductive performance, the effect of Dalmazin (PGF2α) on synchronization, the effect of GnRH and hCG treatment given on the mating day on pregnancy rate, and embryonic mortality after plasma hormone concentrations and embryo development to improve litter size. Male lambs exhibited significantly higher (P<0.05) birth (3.40 ± 0.16 kg) and adult weights (61.67 ± 0.95 kg) compared to females (3.00 ± 0.00 kg and 39.67 ± 0.41 kg, respectively), along with faster growth and higher turnover rates. Seasonal variations showed superior pregnancy rates in spring (62.83%) but higher reproductive efficiency and lower mortality in autumn. In a second experiment, 30 ewes divided into hCG, GnRH, and control groups. hCG and GnRH groups were mated to ram at synchronized estrus by two injections of 2ml PGF2α analogue (Dalmazin) given at 11 days apart. The control group received only saline. The blood samples were collected from jugular venipuncture (3ml) with a disposable syringe from day 2 to 16 after treatment. Plasma progesterone concentrations detected by ELISA were higher (P<0.05) in sheep as compared with saline treated controls and improved reproductive outcomes. The hCG group showed the best performance with a litter size of 1.28%, no embryonic mortality, and increased twinning and fecundity rates. The GnRH group also demonstrated enhanced reproductive efficiency with a litter size of 1.25% and no embryonic loss. The control group had the lowest reproductive outcomes, including a higher embryonic mortality rate. Overall, hCG proved more effective than GnRH in enhancing progesterone levels, fecundity, and prolificacy in Buchi sheep.

## Introduction

Buchi is a dominant sheep breed from a hot region of Pakistan, distinguished by distinct breed features. It can last 72 hours without drinking water. Buchi sheep ranks fourth, following the Kajli, Lohi, and Thalli sheep breeds in Punjab [1]. Significant embryonic mortality has been documented in all farm mammals so far. Maternal recognition of pregnancy, occurring about day 13 post-mating, demands strong signaling among the growing embryo, endometrium, and ovary. Gonadotrophin supplementation during early pregnancy can reduce embryonic losses. GnRH/hCG therapy has been demonstrated to enhance pregnancy rates and litter sizes in sheep [2]. Furthermore, GnRH/hCG administration enhanced conceptus growth, development and implantation, This study will enhance the socio-economic conditions of the population reliant on sheep for their livelihood. An increase in conception rates and litter sizes would also enhance the supply of meat, potentially resulting in lower or stable prices [3]. Embryonic loss is a primary factor constraining reproductive yield in livestock, with reports indicating that the greatest incidence of pregnancy loss occurs in several mammalian species during the early stages of gestation [4].

Reproductive performance throughout various sheep production systems is a fundamental aspect of production units (meat, wool, milk) [5]. Pre-implantation embryonic loss is the major factor limiting optimum reproductive performance in farm animals. In sheep, 30–40% of fertilized eggs are lost during the first 3 weeks of pregnancy [6]. The primary causes of embryonic mortality in sheep include deficiency of luteal function, embryonic-uterine asynchrony, and insufficient embryo development, evidenced by reduced embryo signaling for pregnancy establishment and a reduced autotrophic impact [7]. Insufficient luteal function is a major cause of embryonic losses in sheep. Since the 1980s, many treatment techniques have been employed to enhance luteal function, decrease embryonic mortality, and improve reproductive performance. Luteotropic hormones, including gonadotrophin-releasing hormone (GnRH) and human chorionic gonadotrophin (hCG), were utilized for this aim during the early luteal phase or late luteal phase [8].

The hormones supplied during the luteal phase exert direct or indirect effects on the ovary, leading to the creation of an accessory corpus luteum and an elevation in serum progesterone (P4) levels [9]. Numerous studies indicate that elevated progesterone levels enhance embryonic survival and decrease embryonic losses in ruminants [10]. However, experimental results are inconsistent, conflicting, and frequently inconclusive. Some research indicated that the administration of GnRH or hCG following artificial insemination (AI) or natural mating enhances pregnancy rates and promotes embryonic growth [11]. Moreover, the treatment of hCG or GnRH during the luteal phase reduces embryonic losses and enhances litter size and birth weight in lambs [8].

The primary objective of this study was to determine the optimal timing for administering GnRH and hCG post-mating concerning blood progesterone levels and other reproductive activity in ewes, which was not previously described. The second objective was to compare GnRH with hCG administered on various days of the estrous cycle in ewes superovulated with eCG throughout the spring season [2]. Multiple studies have used hCG to elevate plasma progesterone levels, as this hormone functions similarly to LH. Thus, ewes have utilized it to enhance lambing rates, increase litter size, and promote fetal growth [12]. Administering hCG on day 12 post-mating in cyclic ewes enhances plasma progesterone and oestradiol levels, increases corpora lutea weight, promotes conceptus growth, and improves placental attachment. However, the documented beneficial effects of hCG treatment in adult ewes decreased in cyclic ewe lambs [3].

Anestrus lactating ewes 20–30 days postpartum and anestrus ewes exhibiting low fertility rates enhanced fertility with hCG treatment [13]. This indicates that in sheep groups experiencing challenges in maintaining pregnancy, the injection of hCG may enhance reproductive performance. In prepubertal sheep lambs undergoing estrus induction treatment, the administration of hCG during the luteal phase enhanced the luteal area and plasma progesterone concentration [14]. This study aimed to examine the impact of administering hCG on the day of mating or day 12 post-mating on luteal function, embryo viability, placentation, and reproductive performance in sheep.

Gonadotrophin supplementation may decrease embryonic losses. A solitary administration of the GnRH analog, buserlin, on day 12 of gestation may reduce embryo mortality in cattle and sheep [2]. Similar results have been documented following a solitary administration of human Chorionic gonadotrophin on day 11 or day 12 post-estrus in sheep [2]. The administration of GnRH on days 10, 11, 12, or 13 post-mating has been demonstrated to enhance early embryo survival and pregnancy rates in sheep. The timing of such hormonal therapy may also appear to be significant. It is currently unclear whether GnRH and hCG exert analogous effects on fetal growth and embryo viability. The current study aimed to examine the effects of GnRH and hCG therapies on plasma hormone levels, conceptus development, placentation, and reproductive performance in sheep. GnRH or hCG administration enhances pregnancy rates and reduces embryonic mortality in Buchi sheep [15].

## Materials and Methods

### Experiment No. 1

The study was conducted at the Government livestock farm, Jugaitpeer, district Bahawalpur. The experimental area is located at 72.22 longitude and 29.61 latitude. The region’s native flora mainly consists of drought-tolerant grasses, shrubs, and trees.

#### Stock selection

All ewes and rams on the farm were registered, and data records were acquired for each flock monthly from diverse flocks in the Cholistan Desert, Punjab, Pakistan. Data was collected monthly through first-hand observation of animals. The flocks were chosen to assess the genetic potential, litter size, embryonic mortality, and reproductive performance of Buchi sheep. The sheep were maintained mostly on grazing, along with supplemental feeding of an active ration, and an organized breeding program was implemented on this farm. The newborn lambs were entirely nourished by their mother’s milk for the initial month of life. From the second month onward, they grazed separately from the adults on natural vegetation, allowing them to suckle from their mothers in the morning and evening until the weaning age.

#### Adult body weight

Adult male and female animals were weighed monthly from the beginning of data collection until one year later. Several animals initially chosen for the experiments died, were auctioned, or were slaughtered during the study period and were consequently removed from the final analysis. The animals were assigned a field number via a metallic disc fixed to the neck or through ear tattooing. Each animal was weighed roughly on the same day across various calendar months of the study year, and a record was kept. The overall pregnancy status of each female was recorded through direct observation and information provided by the farmers. The data regarding sex, breed, age, and birth type (twin/single) were recorded for each animal individually; however, the effects of birth type and timing were excluded from the final analysis due to data limitations. The monthly mean weight was later assessed for several groups of animal flocks using standard statistical methods to analyze different aspects.

#### Growth study

A varying number of newborn lambs were selected, with each individual assigned a number and tagged with a metallic disk around the neck. The birth time data was obtained from the specific farmer or through indirect sources accessible to the workers, together with maintained records of sex, kind of birth, and probable breed for each individual separately. The information regarding the type of birth was considered inadequate for a more comprehensive analysis and was therefore skipped from the final data analysis.

Each individual was weighed monthly on a consistent date within the calendar month. Data on the previously born individual was also gathered to clarify the growth rate. Data pooling across multiple age groups was conducted without difficulty because of the comparable growth patterns observed among many individuals within the same stocks. The data on birth weights were verified by existing literature concerning these breeds in the region. The mean weight for each month for each stock was determined from the aggregated data available for various months of age utilizing standard statistical methods [16], with the initial weight recorded considered as the first month of age.

The growth rates were estimated based on the assumption that each month consists of 30 days, and the turnover weight was determined by the daily weight gain of each animal. The calculation involved a standard mathematical conversion, dividing the total average weight gain over various months by 30, although the growth rate was thought to be influenced by differing calendar months and seasonal variations related to the local vegetation supply.

#### Breeding performance

The breeding efficiency of Buchi sheep was assessed by comparing the number of reproductively active females to those that successfully lambed during the study. A selected group of pregnant females was closely monitored to document lambing outcomes and determine annual and biannual lambing patterns. Data on the number of offspring per female post-term was used to calculate the proportions of single, twin, and triplet births, expressed as percentages of total pregnancies.

Annual lamb production per female was calculated by dividing the total number of lambs by the number of observed females. Birth records were analyzed to examine the distribution of births across different months, and the data were grouped to show seasonal trends, specifically the proportions of births in winter and summer. These patterns were then correlated with the area’s vegetation cycle to explore environmental influences on lambing.

Additional information, including the age of male and female sheep (obtained from farmers) and the outcomes of newborns, was used to infer the reproductive age of females. However, data on male reproductive age was limited, as most were sold early, with only a few retained for 2 to 5 years within flocks.

### Experiment No.2

#### Effects of hCG and GnRH on Embryonic Mortality and Reproductive Performance of Buchi Sheep in the Spring Season

##### Animal selection

This study aimed to assess the impact of GnRH or hCG therapy on embryonic mortality and reproductive performance in Buchi sheep throughout the spring season. A total of (n=30) healthy sheep, aged 2-4 years and weighing around 40-45 kg, were utilized for the experiment. Ultrasonography examination was performed on all chosen non-pregnant sheep. All Buchi sheep had unrestricted access to water, shade, and trace mineral salt blocks, and were provided with seasonal fodder and tree foliage.

##### Ultrasonography examination

A real-time, B-mode ultrasound scanner, DUS 60 Digital Ultrasonic Diagnostic Imaging System, with 5-7.5 MHz endocavity linear-array endorectal probes (Edan Endorectal Transducer E741-2) was utilized to differentiate between pregnant and nonpregnant sheep. The scanning protocol described by [17] was employed. After 27 days, ultrasonography was performed again to check embryonic mortality and to confirm pregnancy.

##### Group selection

Randomly allocated the animals (n=30) into three separate treatment groups. This minimized bias and ensured that the groups were equal at the onset of the trial. Group A (n=10) was the Control Group: Sheep that got no intervention. Group B (n=10) was the hCG treatment group, and Group C (n=10) was the GnRH group. All the sheep in various groups were marked with distinct colored ribbons.

Ewes were categorized into three groups. Group A was given no treatment and served as a control. Animals of Groups B and C were first administered a prostaglandin injection of 2ml Dalmazin (PGF2_α_) (Fatro Pharmaceutical Veterinary Industry, Italy) at the onset of the breeding season, and the second injection was administered after 11 days.

##### Estrus synchronization and mating management

Initially, the animals were weighed and assigned to the required groups. Estrus was synchronized in group B and C animals with two intramuscular injections of Dalmazin (PGF2α) prostaglandin analog given 11 days apart. This was validated by placing these females with fertile males in a confined space for natural mating at a ratio of 1:10 (1 male to 10 females). natural mating was considered Day 0 for calculating the gestational age.

##### Treatment protocols and post-treatment monitoring

The sheep were categorized into three groups. Group A (n=10) served as the control with no treatment, while the other functioned as the experimental group. After the second injection of prostaglandin, ewes in estrus were mated naturally. Group B (n=10) received 1ml hCG per sheep, whereas Group C (n=10) animals received 2ml Dalmaralin (GnRH) (Dalmaralin, Fatro Pharmaceutical Veterinary Industry, Italy) per sheep on the day of mating. Ewes returning to estrus after 21 days were observed to determine non-return rates. During lambing, the number and weight of each lamb were recorded [18]. Stillbirths of lambs were recorded for data analysis.

##### Blood sampling

For the progesterone assay, blood samples were obtained via jugular vein puncture from day 2 to 16 following hCG and GnRH injections, utilizing a 5 ml plastic syringe (Lahore Medical Instruments, Pakistan) and a 23-gauge x 1 needle, and subsequently transferred into pre-heparinized 5 ml glass tubes. In each test tube, 1 ml of heparin solution was added to 3 ml of the blood sample. Samples (about 3-5 ml) were collected at 8:00 AM before feeding. Blood samples were collected more often. Blood samples were centrifuged at 3000 rpm for 15 minutes, and the separated serum was placed into Eppendorf tubes and kept at −20 °C until analyzed for progesterone using ELISA. The hemolyzed samples were eliminated.

Graphical methodology of the present study (Exp 2) is shown in Fig.1.

**Fig. 1:**
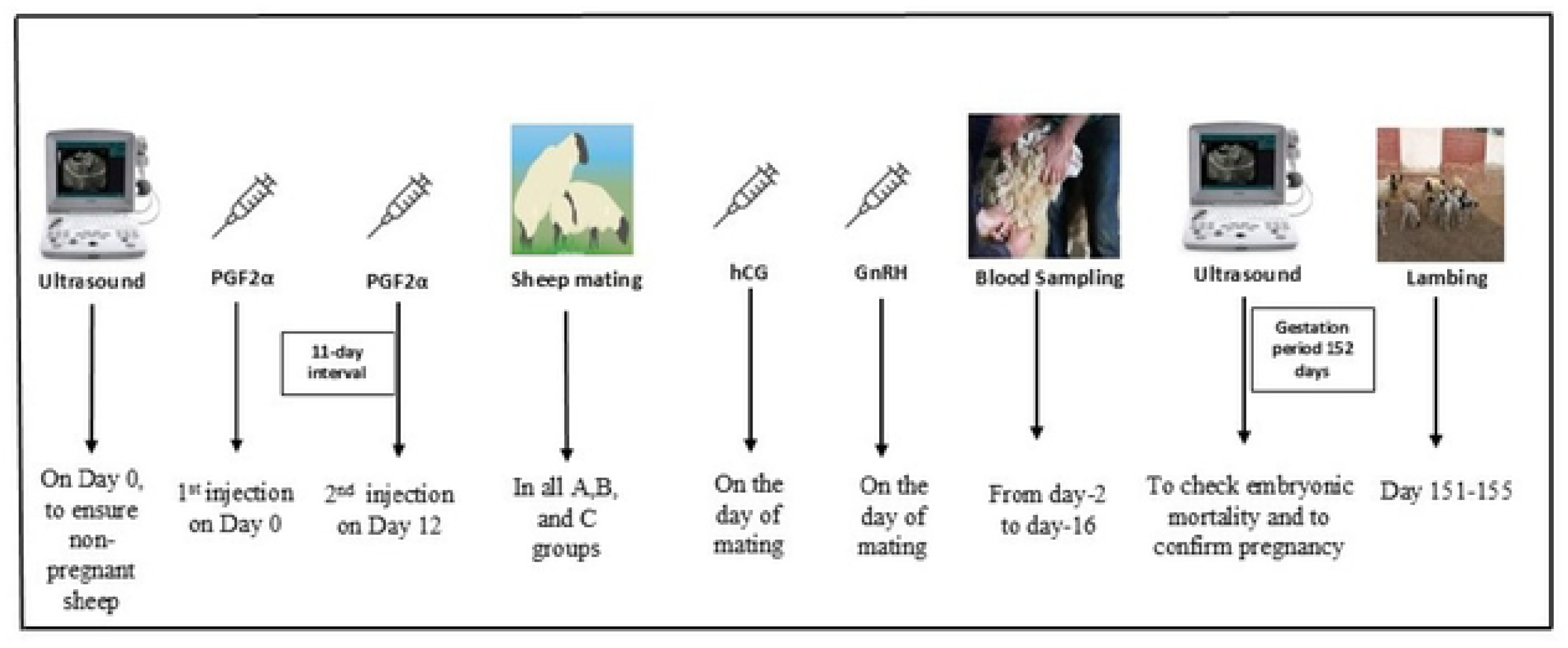
Graphical methodology of the present study (Exp 2)

## Results

### Adult and lambs birth body weight

Buchi sheep’s mean (± SEM) adult and lambs’ birth body weight of experimental animals is shown in (Table 1). A significant difference (P<0.05) was observed in the adult body weight and lambs birth weight between males and females.

**Table 1:** Mean ± SEM adult body weight (kg) and birth body weight of Buchi lambs.

| Gender | Body weight (kg)<br>(Adult) | Birth body weight (kg)<br>(Lambs) |
| --- | --- | --- |
| Male | 61.67 ± 0.96<sup>a</sup> | 3.40 ± 0.16<sup>a</sup> |
| Female | 39.67 ± 0.41<sup>b</sup> | 3.00 ± 0.00<sup>b</sup> |

Values sharing different superscripts in a column differed significantly ( $P < 0.05$ )

### Seasonal variation in adult body weight

Males exhibited greater seasonal body weight variability, while females maintained a steady growth trend, with significant (P<0.05) weight differences observed between the sexes due to environmental and nutritional factors in (Table 2)

**Table 2:** Seasonal Variation in Mean ± SEM adult body weight of Buchi sheep.

| (Mean ± SEM) Body Weight (Kg) | | |
| --- | --- | --- |
| Age (Months) | Male | Female |
| 1 | 54.25 ± 1.55<sup>a</sup> | 39.03 ± 0.44<sup>b</sup> |
| 2 | 57.75 ± 1.49<sup>a</sup> | 39.37 ± 0.36<sup>b</sup> |
| 3 | 62.00 ± 0.58<sup>a</sup> | 39.22 ± 0.33<sup>b</sup> |
| 4 | 63.00 ± 0.41<sup>a</sup> | 39.70 ± 0.28<sup>b</sup> |
| 5 | 60.50 ± 0.29<sup>a</sup> | 39.36 ± 0.48<sup>b</sup> |
| 6 | 64.25 ± 0.85<sup>a</sup> | 40.05 ± 0.43<sup>b</sup> |
| 7 | 61.25 ± 1.38<sup>a</sup> | 38.39 ± 0.41<sup>b</sup> |
| 8 | 63.00 ± 1.83<sup>a</sup> | 39.16 ± 0.36<sup>b</sup> |
| 9 | 65.00 ± 1.91<sup>a</sup> | 39.16 ± 0.45<sup>b</sup> |
| 10 | 62.50 ± 1.89<sup>a</sup> | 40.91 ± 0.42<sup>b</sup> |
| 11 | 64.25 ± 1.70<sup>a</sup> | 42.04 ± 0.39<sup>b</sup> |
| 12 | 65.75 ± 1.5<sup>a</sup> | 43.47 ± 0.33<sup>b</sup> |

Values sharing different superscripts in a row differed significantly ( $P < 0.05$ )

### The growth rate of Buchi lamb

A 12-month growth study of Buchi lambs revealed consistent weight gain in both sexes, with males growing significantly faster than females (P<0.05) (Table 3). At 1 month, male lambs weighed 5.6 ± 0.11 kg compared to 5.16 ± 0.11 kg in females. This gap widened over time, with males reaching 42.52 ± 0.91 kg and females 28.79 ± 0.68 kg by 12 months. The most pronounced growth in males occurred between the 9th and 12th months. These findings reflect typical sexual dimorphism in sheep, with males achieving higher body weights. The small SEM values indicate uniform growth trends across the population.

**Table 3:** Mean ± SEM Growth rate of Buchi lamb.

| (Mean ± SEM) Body weight (kg) | | |
| --- | --- | --- |
| Age (Months) | Male | Female |
| 1 | $5.6 \pm 0.11^a$ | $5.16 \pm 0.11^b$ |
| 2 | $8.7 \pm 0.15^a$ | $7.88 \pm 0.13^b$ |
| 3 | $12.59 \pm 0.21^a$ | $11.06 \pm 0.19^b$ |
| 4 | $17.27 \pm 0.24^a$ | $15.33 \pm 0.34^b$ |
| 5 | $20.89 \pm 0.32^a$ | $19.91 \pm 0.37^b$ |
| 6 | $23.31 \pm 0.49^a$ | $21.68 \pm 0.41^b$ |
| 7 | $24.98 \pm 0.69^a$ | $20.87 \pm 0.53^b$ |
| 8 | $28.04 \pm 0.79^a$ | $22.00 \pm 0.62^b$ |
| 9 | $31.17 \pm 0.92^a$ | $23.12 \pm 0.56^b$ |
| 10 | $35.08 \pm 1.06^a$ | $24.74 \pm 0.62^b$ |
| 11 | $39.48 \pm 0.98^a$ | $26.54 \pm 0.58^b$ |
| 12 | $42.52 \pm 0.91^a$ | $28.79 \pm 0.68^b$ |

Values sharing different superscripts in a row differed significantly ( $P < 0.05$ )

### Body weight (kg) and turnover weight(g) of male and female Buchi lambs

A 12-month study assessed the body and turnover weights of male and female Buchi lambs (Table 4). Both sexes showed steady growth, but males consistently outperformed females in both parameters. Male lambs increased from 5.24 ± 0.13 kg to 42.52 ± 0.91 kg in body weight and from 173.03 ± 4.10 g to 1412.6 ± 32.7 g in turnover weight. Females grew from 5.38 ± 0.17 kg to 28.79 ± 0.68 kg, and their turnover weight rose from 177.14 ± 5.74 g to 929.1 ± 54.9 g. While no significant differences were noted during the first five months (P>0.05), a significant difference (P<0.05) emerged from months 6 to 12, with males showing accelerated growth. The data confirms typical sexual dimorphism in sheep, with males exhibiting greater and more rapid development, likely influenced by genetic and hormonal factors.

**Table 4:**
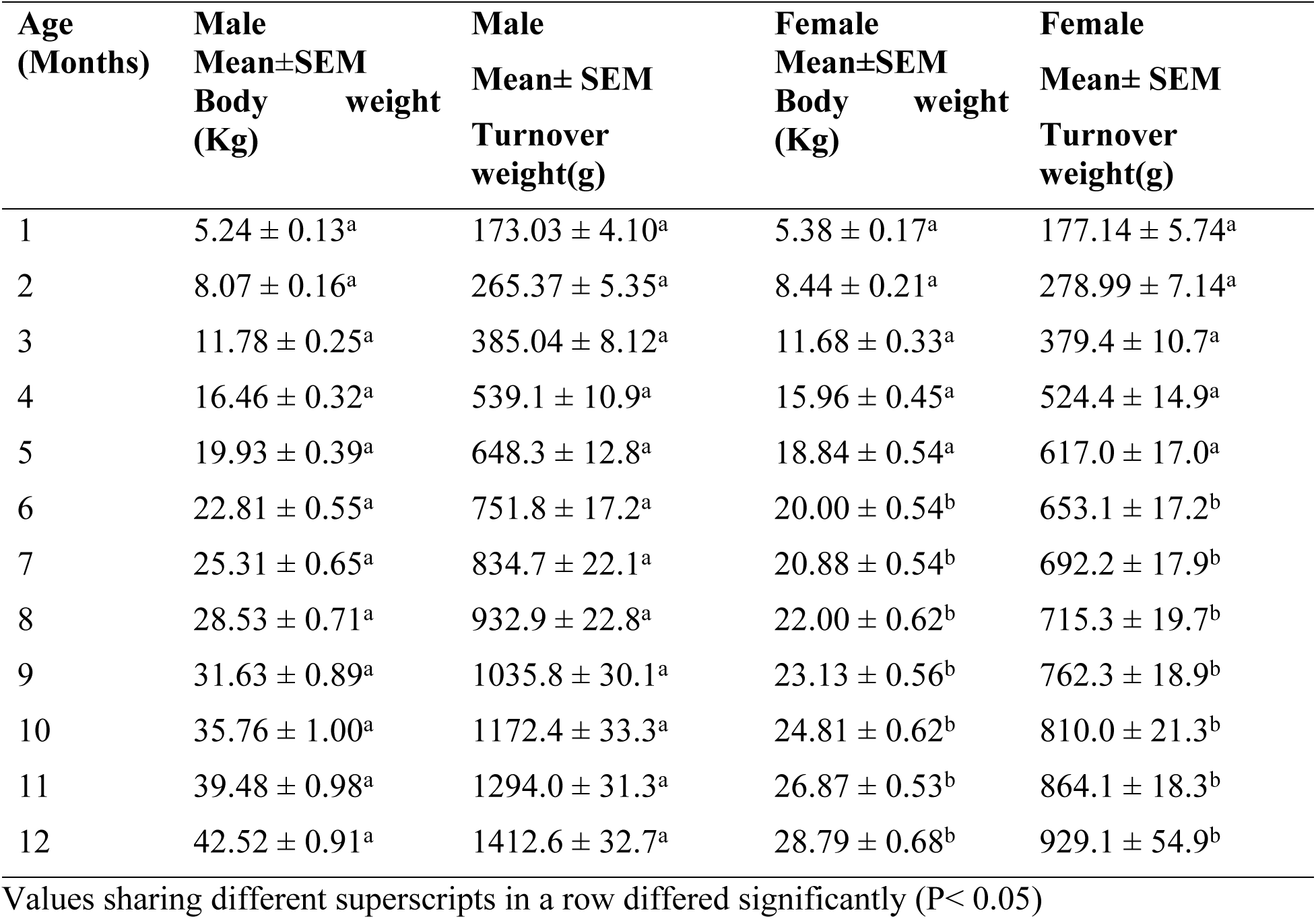
Mean ± SEM of body weight (kg) and Turnover weight (g) of male and female Buchi lambs.

| Age<br>(Months) | Male<br>Mean±SEM<br>Body weight<br>(Kg) | Male<br>Mean± SEM<br>Turnover<br>weight(g) | Female<br>Mean±SEM<br>Body weight<br>(Kg) | Female<br>Mean± SEM<br>Turnover<br>weight(g) |
| --- | --- | --- | --- | --- |
| 1 | $5.24 \pm 0.13^a$ | $173.03 \pm 4.10^a$ | $5.38 \pm 0.17^a$ | $177.14 \pm 5.74^a$ |
| 2 | $8.07 \pm 0.16^a$ | $265.37 \pm 5.35^a$ | $8.44 \pm 0.21^a$ | $278.99 \pm 7.14^a$ |
| 3 | $11.78 \pm 0.25^a$ | $385.04 \pm 8.12^a$ | $11.68 \pm 0.33^a$ | $379.4 \pm 10.7^a$ |
| 4 | $16.46 \pm 0.32^a$ | $539.1 \pm 10.9^a$ | $15.96 \pm 0.45^a$ | $524.4 \pm 14.9^a$ |
| 5 | $19.93 \pm 0.39^a$ | $648.3 \pm 12.8^a$ | $18.84 \pm 0.54^a$ | $617.0 \pm 17.0^a$ |
| 6 | $22.81 \pm 0.55^a$ | $751.8 \pm 17.2^a$ | $20.00 \pm 0.54^b$ | $653.1 \pm 17.2^b$ |
| 7 | $25.31 \pm 0.65^a$ | $834.7 \pm 22.1^a$ | $20.88 \pm 0.54^b$ | $692.2 \pm 17.9^b$ |
| 8 | $28.53 \pm 0.71^a$ | $932.9 \pm 22.8^a$ | $22.00 \pm 0.62^b$ | $715.3 \pm 19.7^b$ |
| 9 | $31.63 \pm 0.89^a$ | $1035.8 \pm 30.1^a$ | $23.13 \pm 0.56^b$ | $762.3 \pm 18.9^b$ |
| 10 | $35.76 \pm 1.00^a$ | $1172.4 \pm 33.3^a$ | $24.81 \pm 0.62^b$ | $810.0 \pm 21.3^b$ |
| 11 | $39.48 \pm 0.98^a$ | $1294.0 \pm 31.3^a$ | $26.87 \pm 0.53^b$ | $864.1 \pm 18.3^b$ |
| 12 | $42.52 \pm 0.91^a$ | $1412.6 \pm 32.7^a$ | $28.79 \pm 0.68^b$ | $929.1 \pm 54.9^b$ |

Values sharing different superscripts in a row differed significantly ( $P < 0.05$ )

### Reproductive performance of Buchi sheep belonging to different seasons

A seasonal comparison of Buchi sheep breeding performance (Table 5), revealed notable differences between spring and autumn. In spring, 62.83% of 113 sheep were pregnant, producing 62 lambs with a lambing rate of 87.32%. Single births dominated (93.54%), while twin births were rare (6.45%), and abortion/dead birth cases were relatively higher (14.5%). In contrast, autumn showed a lower pregnancy rate (42.5% of 120 sheep) but a higher lambing rate (92.15%) with fewer abortion cases (2.82%). Single births made up 87.23%, and twin births increased slightly to 12.76%. These variations suggest that seasonal environmental factors, nutrition, and management practices influence reproductive outcomes in Buchi sheep.

**Table 5:** Mean ± SEM Reproductive performance of Buchi sheep belonging to different seasons.

| Parameters | Spring | % | Autumn | % |
| --- | --- | --- | --- | --- |
| No. of sheep | 113 |  | 120 |  |
| Lambing pattern |  |  |  |  |
| Pregnant Once/year | 71 | 62.83 | 51 | 42.5 |
| Non-Pregnant | 42 | 37.17 | 69 | 57.5 |
| Litter size. |  |  |  |  |
| Single birth (%) | 58 | 93.54 | 41 | 87.23 |
| Twins birth (%) | 2*(4) | 6.45 | 3*(6) | 12.76 |
| Triplet birth (%) | 0 |  | 0 |  |
| No. of lambs. |  |  |  |  |
| Lambs born /year | 62 | 87.32 | 47 | 92.15 |
| Dead birth (%) | 9 | 14.5 | 6 | 12.76 |
| Aborted | 02 | 2.82 | 01 | 1.96 |

\*The total number of lambs produced by 2 sheep was 4 in the spring.
\*Total number of lambs produced by 3 sheep was 6 in autumn.

### Embryonic mortality and reproductive performance of Buchi sheep injected with saline (control), hCG and GnRH during spring season

The comparison of embryonic mortality and reproductive performance in Buchi sheep treated with saline (control), human chorionic gonadotropin (hCG) and Gonadotropin-releasing hormone (GnRH) during spring season (Table 6), highlights the positive impact of hCG and GnRH. There was a non-significant difference (P>0.05) in gestation length of all groups. Birth weights were also significantly higher (P<0.05) in the hCG group, and no dead births occurred in hCG and GnRH. These results suggest that hCG and GnRH treatment significantly enhances reproductive performance, increases litter size, and reduces embryonic mortality in Buchi sheep during the spring season. But the GnRH is more effective.

**Table 6:**
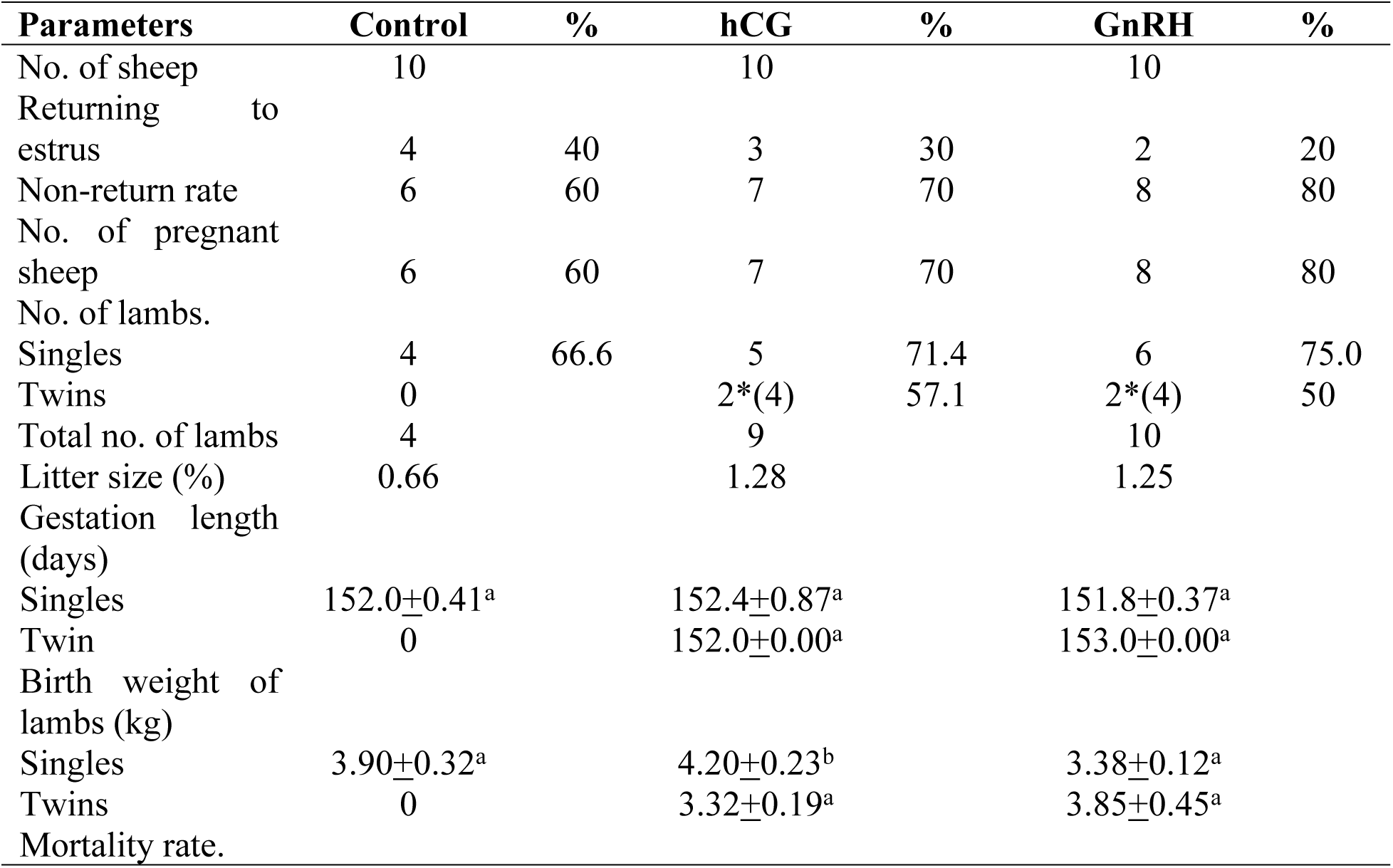
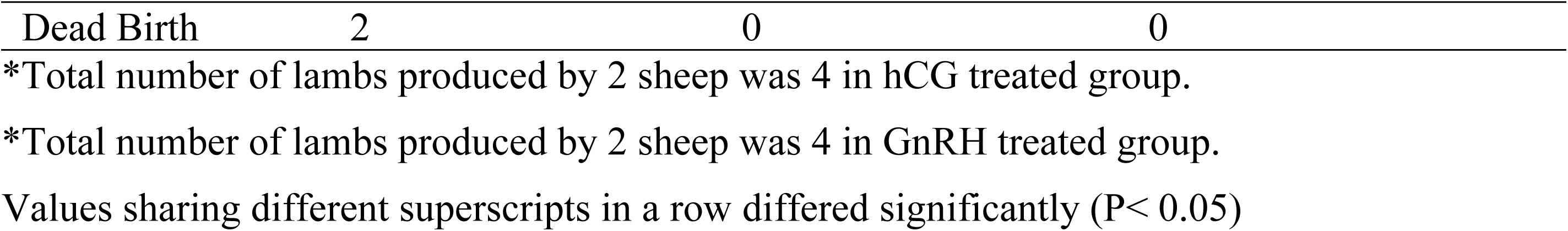
Embryonic mortality and reproductive performance of Buchi sheep injected with saline (control), hCG and GnRH in the spring season in different groups at Govt livestock farm, Jugait peer District Bahawalpur.

### Progesterone concentration levels in the spring season

Table 7 summarizes the mean (± SEM) progesterone concentrations in Buchi sheep across different days (2, 5, 8, 11, and 16) for Control, hCG, and GnRH-treated groups. Initially, on day 2, the GnRH group showed significantly lower progesterone levels (2.58 ± 0.10) compared to the control (9.29 ± 2.73) and hCG group (10.07 ± 2.99). However, from day 5 onward, all groups exhibited rising progesterone concentrations, with the hCG group consistently showing the highest levels across all time points. By day 16, the hCG group reached a peak progesterone level of 28.05 ± 1.99, followed by GnRH (26.41 ± 4.10), and the control group (20.64 ± 4.02). These findings indicate that both hCG and GnRH treatments enhance progesterone production over time, with hCG having the most pronounced effect. This suggests a stronger luteotrophic influence of hCG, potentially supporting improved reproductive outcomes in treated animals. Plasma progesterone levels in Buchi sheep during the spring season showed in Fig. 2.

**Fig. 2:**
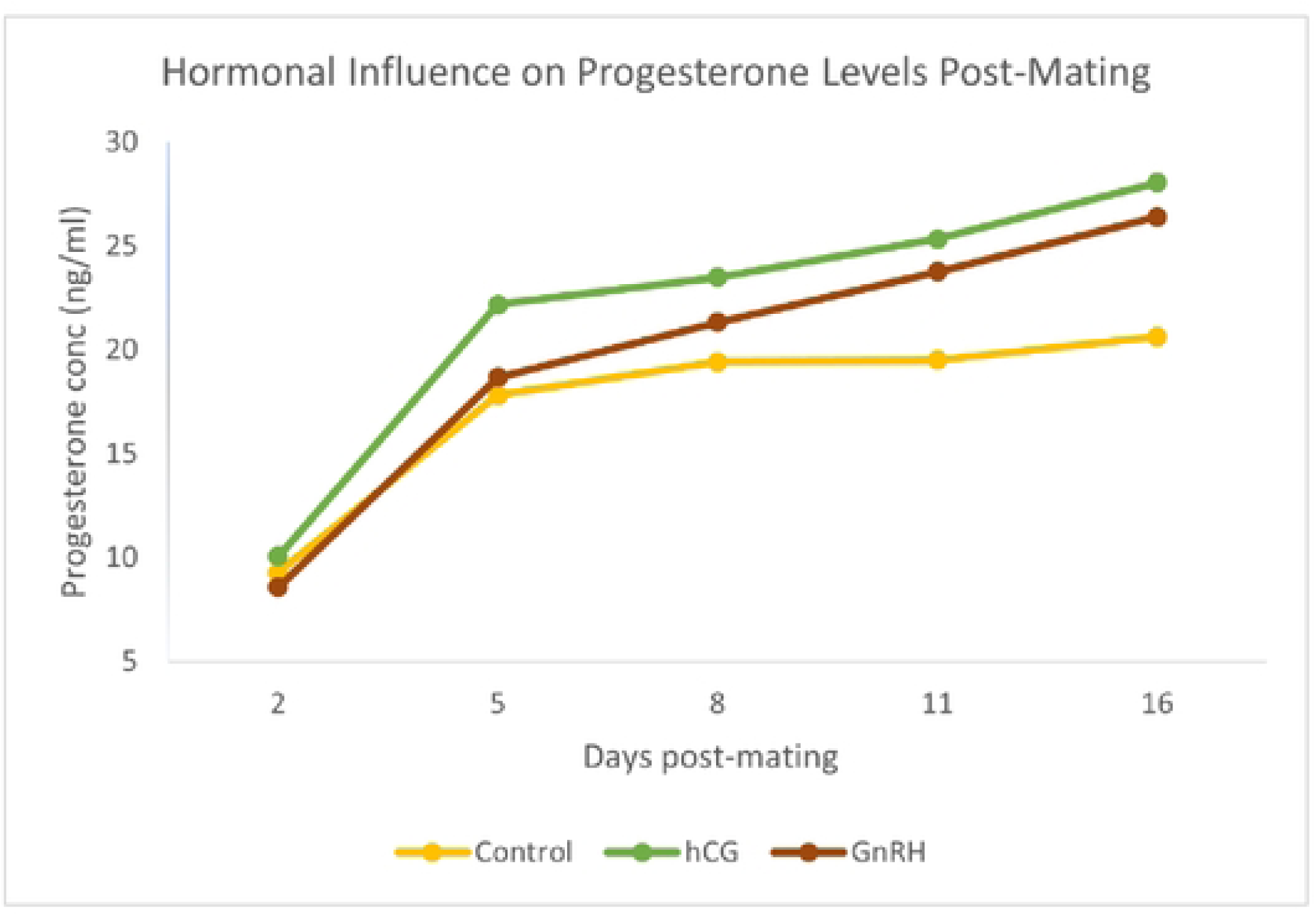
Plasma progesterone levels in Buchi sheep during the spring season.

**Table 7:** Mean ±SEM Plasma progesterone levels in Buchi sheep during the spring season.

Days of sampling
| Hormones | 2 | 5 | 8 | 11 | 16 |
| --- | --- | --- | --- | --- | --- |
| Control | $9.29 \pm 2.73$ | $17.83 \pm 3.16$ | $19.42 \pm 2.33$ | $19.51 \pm 1.76$ | $20.64 \pm 4.02$ |
| hCG | $10.07 \pm 2.99$ | $22.19 \pm 4.01$ | $23.50 \pm 3.20$ | $25.33 \pm 3.64$ | $28.05 \pm 1.99$ |
| GnRH | $8.58 \pm 0.10$ | $18.67 \pm 1.05$ | $21.34 \pm 4.58$ | $23.76 \pm 0.73$ | $26.41 \pm 4.10$ |

## Discussion

### Adult and lamb birth body weight

The Mean ± SEM adult body weight of males and females recorded during the study period was 61.67 ± 0.95 kg and 39.67 ± 0.41 kg, respectively. A significant difference (P<0.05) was observed in the adult body weight between males and females. The previous study showed that the adult body weight of male and female sheep in Pakistan has revealed significant variation in different breeds. The Hissardale breed exhibits adult weights of 66.4 kg for males and 41.94 kg for females, while the Awassi breed shows slightly higher weights at 67.89 kg for males and 45.34 kg for females [19]. The Lohi breed presents a mean adult weight of 61.75 kg for rams and 54.26 kg for ewes [20]. On the other hand, the Kacchi breed has lower body weights, with males averaging 51.24 kg and females 35.19 kg [21]. These findings highlight the diversity in sheep body weight between different breeds in Pakistan, influenced by genetic and environmental factors. The Balochi and Beverigh breeds also reflect this trend, with male weights at 39.18 kg and 35.80 kg for females at 24 months [22]. These findings indicate a diverse range of body weights influenced by breed and management practices, highlighting the importance of genetic potential in sheep production in Pakistan. Rams consistently exhibit greater body weight than female sheep (ewes) across various ages and breeds, indicating a strong genetic basis for sexual dimorphism in body weight [23]. Early nutritional conditions significantly impact adult body composition, with undernutrition during critical developmental periods leading to sex-specific metabolic changes. Males show increased fat mass and altered lipid metabolism compared to females [24,25]. Environmental factors, such as seasonal conditions and management practices, also play a crucial role in liveweight variation, affecting both sexes but with differing impacts on weight loss and gain [26]. The gender of the newborn lamb significantly affects birth weight, with male lambs (3.40 ± 0.16) being significantly (P<0.05) heavier than female lambs (3.00 ± 0.00).

Similarly, [27] documented that Thalli sheep lambs born in the spring season were marginally heavier (4.05±0.12 kg) than those born in the fall season (4.01±0.06 kg), while single lambs (4.24±0.00 kg) were shown to be heavier than twin lambs (3.68±0.01 kg). The male lambs exhibited greater weight than the females, as indicated. The average birth weights for male and female lambs were 4.21±0.10 kg and 3.85±0.08 kg, respectively. [28] noted that season of birth and sex significantly influenced birth weight. The birth weight of lambs born in spring was much greater than that of lambs born in October, a fact also corroborated by the present study. The outcomes of various surroundings. The average birth weight of Hissardale male and female lambs was 4.72±0.80 kg and 4.46±0.84 kg, respectively; likewise, the birth weight of Awassi male and female lambs was 4.89±0.16 kg and 4.30±0.14 kg, respectively [19]. Lambs born as singles generally exhibit higher birth weights compared to those born as twins, with significant differences noted in certain sheep genotypes [29]. The decline in birth weight is also associated with increased litter size, indicating a linear relationship between litter size and lamb birth weight [30]. Ewes that receive adequate nutrition throughout gestation tend to produce heavier lambs, emphasizing the importance of maternal diet [31]. Environmental conditions, such as the season and the age of the ewe, also play a crucial role. Lambs born in spring-summer tend to weigh more, while younger and older ewes produce lighter lambs [32].

### Seasonal variation in adult body weight

The average seasonal variation in the (mean ± SEM) body weight of adult male and female Buchi sheep over 12 months. Both sexes showed a consistent pattern of cyclic variation. In males, the body weight fluctuates over the year, starting at 54.25 ± 1.55 kg in the first month and gradually increasing to 65.75 ± 1.49 kg by the 12^th^ month. While the female Buchi sheep maintain a more stable body weight throughout the year. Their weight starts at 39.03 ± 0.44 kg in the first month and gradually increases, reaching 43.47 ± 0.33 kg by the 12^th^ month. Overall, in 12 months, a significant difference (P<0.05) was observed in the body weight of male and female Buchi sheep.

Studies indicate that body weight fluctuates significantly across seasons, with peaks often observed in autumn and declines in spring. For instance, adult ewes of various breeds, including Malpura and Avimaans, showed a mean weight of 31.10±0.067 kg, with the highest weights recorded in October (34.49±0.217 kg) and the lowest in April (27.84±0.218 kg) [33]. Buchi sheep exhibit notable seasonal weight fluctuations influenced by environmental factors, which can differ from other breeds. Their preweaning and postweaning growth traits are significantly affected by year and season, with average weights at weaning adjusted to 120 days being 12.55 kg and 33.78 kg at 365 days [1]. Seasonal variations in adult sheep body weight result from feed availability, climate, reproductive cycles, hormonal changes, and management practices. Weight gain occurs in spring and early summer due to abundant forage, while winter and late summer bring weight loss due to poor feed quality and climatic stress [34].

### The growth rate of Buchi lamb

The lamb’s weight consistently grows until 12 months of age. The average growth rates for males and females during the initial four months of age were 5.6 to 17.27 kg and 5.16 to 15.33 kg, respectively, indicating a significant difference (P ˂ 0.05). This was followed by a gradual increase until the 5th to 8th month, reaching 20.89 to 28.04 kg in males and 19.91 to 22.00 kg in females. Growth continued until the 9th to 12th months, the males increasing from 31.17 to 42.52 kg and females from 23.12 to 28.79 kg, respectively. The males reported more weight than the females across all age groups. The examination of growth rates across various ages for both genders indicates that, while considerable weight differences between male and female lambs at comparable ages, as determined by maximum likelihood approaches, males consistently outweighed females at all ages. The growth rate of Buchi lambs over 12 months exhibits considerable variety in comparison to other sheep breeds. In a study, Balochi lambs attained an average weight of 30.65 ± 1.92 kg for males and 27.42 ± 1.04 kg for females at 12 months [22]. Conversely, Texel lambs demonstrated a wider spectrum of growth rates, with heritabilities for these rates ranging from 0.14 to 0.74 [35]. Individual lambs exhibit significant genetic variability, influencing their growth rates. For instance, lambs from ewes rearing singletons tend to show higher average daily gains compared to those from ewes with twins or triplets [36]. Effective management practices, including monitoring growth rates and adjusting feeding regimens, can enhance productivity and profitability in lamb production [37].

### Body weight (kg) and turnover weight(g) of male and female Buchi lambs

The turnover rate based on the daily weight gain per animal across various ages, sexes, and months. The gender-specific turnover rate was 173,265,385,539 g/day for male lambs and 177,278,379,524 g/day for female lambs during the initial four months of age, respectively. The average turnover rates of lambs, categorized by gender throughout various seasons, showed significant differences (P˂0.05). The overall growth rate of the current stock increased by 16 kg in males and 15 kg in females over the initial four months of age, respectively. The growth rate subsequently increased by 28.53 kg for males and 22.00 kg for females from months 5 to 8, followed by a gradual increase of 42.52 kg for males and 28.79 kg for females in the older age range of 9-12 months. The males showed a superior growth rate compared to the females. The mean (± SEM) body weight and turnover weight (± SEM) of male and female Buchi lambs over 12 months can be compared with other breeds based on various studies. Baluchi male sheep at 12 months had a mean body weight of approximately 35 kg, with growth rates of 222.4 g/day for those in a growth group [38]. In contrast, lambs from other breeds, such as the improved Jezersko-Solcava, had an average body weight of 22.9 kg at weaning, with males showing higher daily gains. Notably, sex differences were significant, with males generally exhibiting better growth performance than females [39]. Males generally exhibit greater body weights than females due to higher muscle mass, Genetic Influences and fat deposition. For instance, male sheep in a study showed significant weight differences, averaging 36.76 kg compared to 32.12 kg for females [40]. Climatic conditions significantly impact body weight, with variations in weather affecting growth rates in different regions. For example, lamb weights were influenced by spring and summer weather conditions across various grazing areas [41].

### Reproductive performance of Buchi sheep belonging to different seasons

Various parameters correlated with the reproductive efficiency of Buchi sheep, classified by gender. In the present study, all the ewes showed a biannual lambing trend. The female fertility rate in spring was 62.83%, whereas in autumn it was 42.5%. A notable variation (P < 0.05) was seen in sheep fertility rates across seasons. Single birth rates were 93.54% in spring and 87.23% in autumn. Twin births were recorded at a rate of 6.45% in spring and 12.76% in autumn, with no occurrences of triplet births in either season. The death rate was 14.5% in spring and 12.76% in autumn, whereas the abortion rate was 2.82% in spring and 1.96% in autumn. Similarly, the female fertility rates for Lohi and Kacchi sheep were 86.3% and 80%, respectively. No significant difference was observed in the reproductive rates of sheep across varying flock sizes for both breeds. The incidence of twin births was 16.34% in Lohi sheep and 13.60% in Kacchi sheep, with no instances of triplet litters recorded in either breed. The average litter size was 1.53 per sheep. Lambing transpired throughout two distinct seasons: August to November and February to April. In conclusion, the Lohi breed exhibits superior genetic reproductive performance, birth weight, and growth potential. The lambs exhibit a superior survival rate. The Lohi sheep exhibit strong breeding potential and are well adapted to thrive in both dry and irrigated conditions of the region [20]. Various studies on stock and reproductive rates in Pakistan indicate significant variability in sheep fertility. The prevalence of sheep in flocks in the current study is significantly higher than previously published figures: Karakuli at 75%, Balali at 69%, and Randozai at 64% by, as well as Horo at 70.1% and Menz at 79.5% by [42]. However, some studies indicate a greater reproduction rate: 96.1% in Lohi, 97.1% in Afghan, and 90% in the general nomadic flock of Lebanon [43]. The variation in reproductive performance of sheep across different seasons is influenced by a combination of environmental factors, hormonal changes, and management practices. Seasonal breeding patterns are primarily dictated by photoperiod, with ewes exhibiting increased reproductive activity as day length decreases, particularly in autumn. This seasonal behavior is further affected by climatic conditions, such as temperature, precipitation, and sunlight, which can impact fertility rates and lambing success (44,45).

### Embryonic mortality and reproductive performance of Buchi sheep injected with saline (control), hCG and GnRH during spring season

The reproductive performance in the hCG and GnRH-treated ewes was significantly higher (P<0.05) than that of the control group, with no embryonic death was seen in the hCG and GnRH treated group compared to the control group. A similar study by [2] showed that hCG treatment improved embryo survival in hCG-treated animals with more triplet, quadruplet, and quintuplet births than the saline-treated controls.

Consistent with our findings, previous studies have indicated that hCG injection in the early stages of pregnancy enhances lamb production and reproductive performance in sheep [2]. [46] asserted that gonadotrophins are essential for follicular development. The dosage and timing of hCG injection are crucial for enhancing luteal activity evaluated the dose- and time-dependent impact of hCG on luteal function in anestrus ewe lambs [14]. The variation in embryonic mortality and reproductive performance between saline-injected (control) and hCG-treated sheep is attributed to hCG’s effectiveness in inducing estrus, promoting follicular growth, and enhancing ovulation rates, leading to improved embryonic implantation rates [47]. For instance, administering hCG shortly after estrus observation can lead to synchronized ovulation and improved pregnancy rates [48]. The physiological state of the ewes, including their reproductive history and health, can influence the effectiveness of hormonal treatments. Ewes treated with hCG showed better follicular development and ovulation rates compared to controls [49].

Similarly, finding was observed by [50], a favorable impact on ovulation synchronization in ewes using GnRH - PGF2α - GnRH during the non-breeding season in a Karakul flock. The effect of exogenous GnRH injection may differ depending on the stage of the estrous cycle in the treated animals [46]. Ewes treated with GnRH or hCG showed improved reproductive metrics, such as higher twinning rates and fecundity compared to control groups receiving saline [48]. GnRH treatment resulted in significantly elevated serum progesterone levels, which are essential for maintaining pregnancy [49]. GnRH treatment altered metabolite levels related to collagen and prostaglandin synthesis, potentially affecting embryo implantation and overall pregnancy success. The study found that GnRH treatment significantly reduced pregnancy rates in Huyang ewes (72.2% vs. 82.9% for saline control, P < 0.05), indicating potential increased embryonic mortality and impaired reproductive performance associated with GnRH administration during artificial insemination [51]. The timing of GnRH administration relative to estrus onset significantly influenced mating success and pregnancy rates, with variations noted across different sheep breeds [52]. Different dosages of GnRH can also lead to varying reproductive outcomes. For example, higher doses of GnRH did not significantly improve ovulation or pregnancy rates compared to control groups, indicating that optimal dosing is crucial for achieving desired reproductive outcomes [48].

### Plasma progesterone levels in Buchi sheep during spring season

The progesterone concentrations (mean ± SEM) across different days (2, 5, 8, 11, and 16) for Control, hCG, and GnRH groups. Initially, the hCG group had slightly higher levels than the control, while the GnRH group had the lowest. Progesterone levels increased in all groups over time, with hCG consistently showing the highest concentrations, followed by GnRH and then the control. By day 16, the hCG group had the highest progesterone concentration, indicating that both hCG and GnRH treatments elevated progesterone levels, with hCG having the most significant effect. The first reason for doing this study was to identify whether different hormonal programs at different times after mating could improve the concentration of progesterone as a pregnancy-retaining hormone in sheep or not. The results of this study showed that the use of GnRH and hCG during different days (days 1, 2, 5, 7 and 12) after mating in Lake-Ghashghaei ewes significantly increased serum progesterone concentration compared to the control group that was consistent with the results of other studies [53,54]. In those studies, despite the time of using these hormones, the concentration of progesterone compared to the control group was increased significantly. hCG has similar activity to Luteinizing Hormone (LH) with a longer half-life than LH and causes luteotropic stimulation in the corpus luteum [55]. These effects can be done either by converting small cells into large corpus luteum or by increasing the size of large luteal cells [54]. Increased progesterone levels can be due to an increase in the number of extra corpus luteum on the surface of the ovary, and or increase the number of luteal cells that secrete progesterone [56]. In some other studies, increasing the pregnancy rate and the rate of lambing was reported when using GnRH and hCG in the time of mating or on different days after mating in different breeds of sheep. Differences in results can be attributed to the different protocols used, management systems, nutritional or physiological status of ewes and experimental conditions [57]. Several previous studies reported that hCG increases the embryo’s size and length, increases the secretion of interferon tau, luteal weight, and increases placental weight [18].

## Acknowledgments

We are grateful the research funding provided through the research supporting project No.3910/ORIC/IUB/2021 by ORIC, the Islamia University of Bahawalpur.

